## Supplementary figures and images for "Signaling by the integrated stress response kinase PKR is fine-tuned by dynamic clustering"

### Supplemental Figure 1

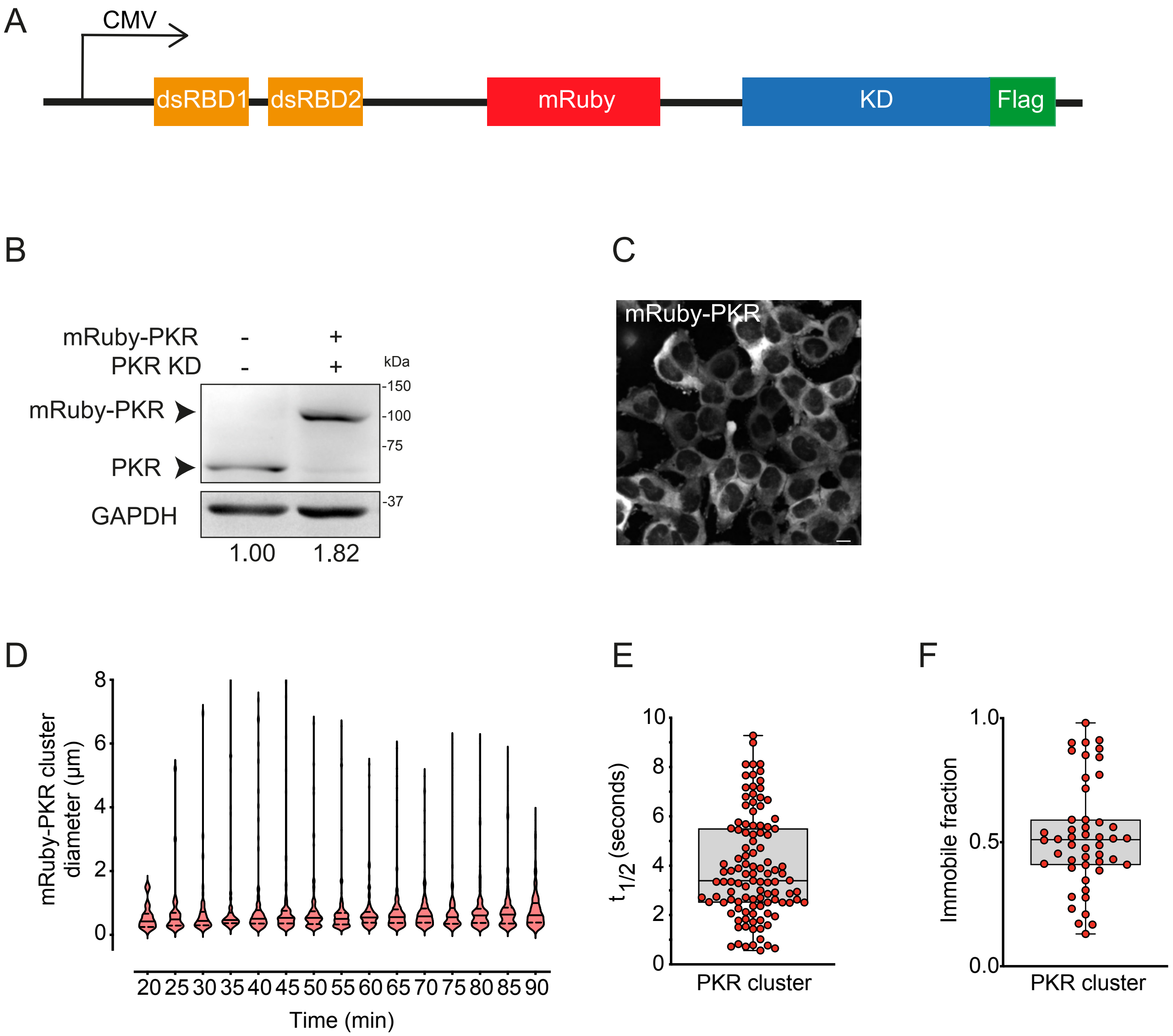

### Supplemental Figure 2.1

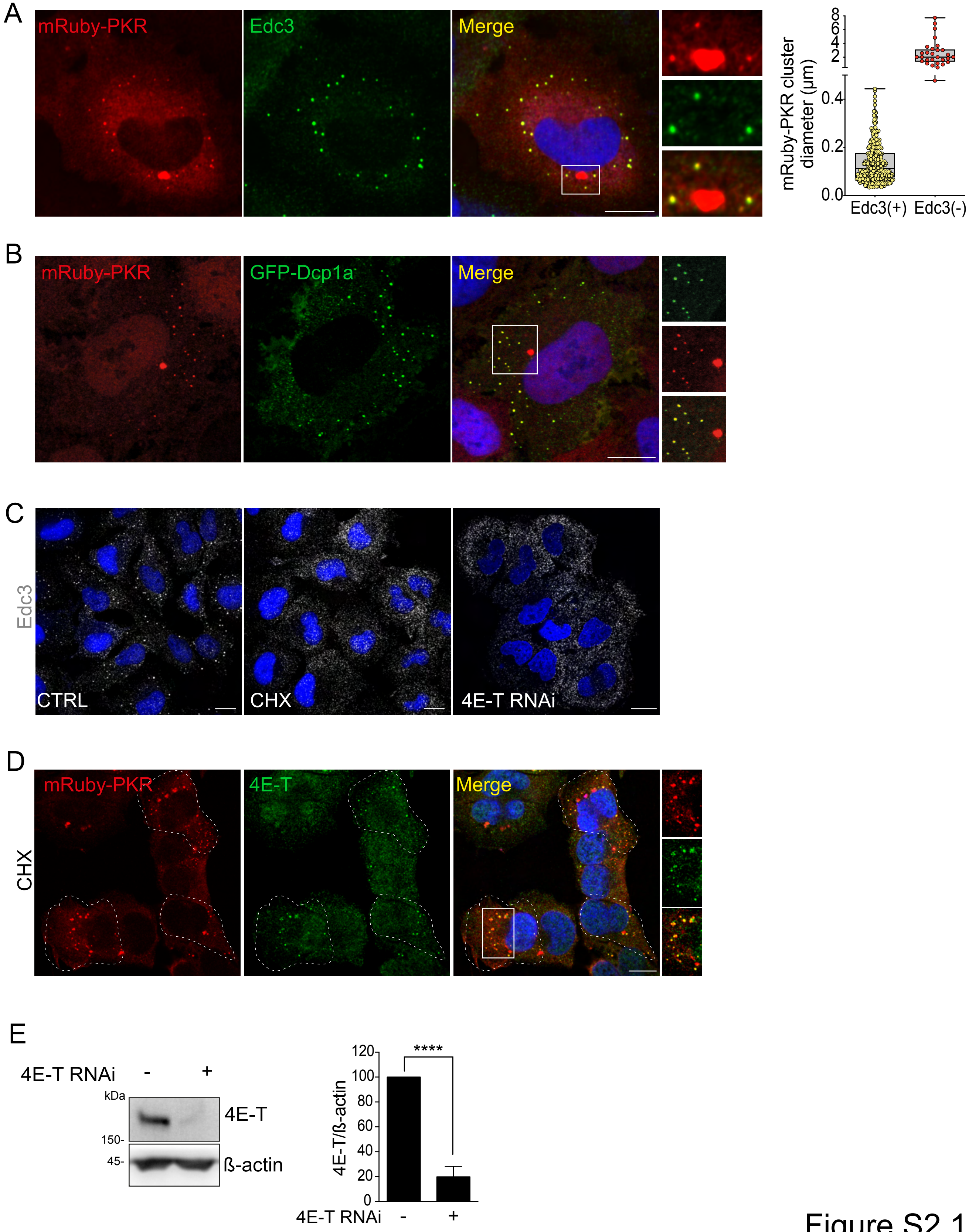

### Supplemental Figure 2.2

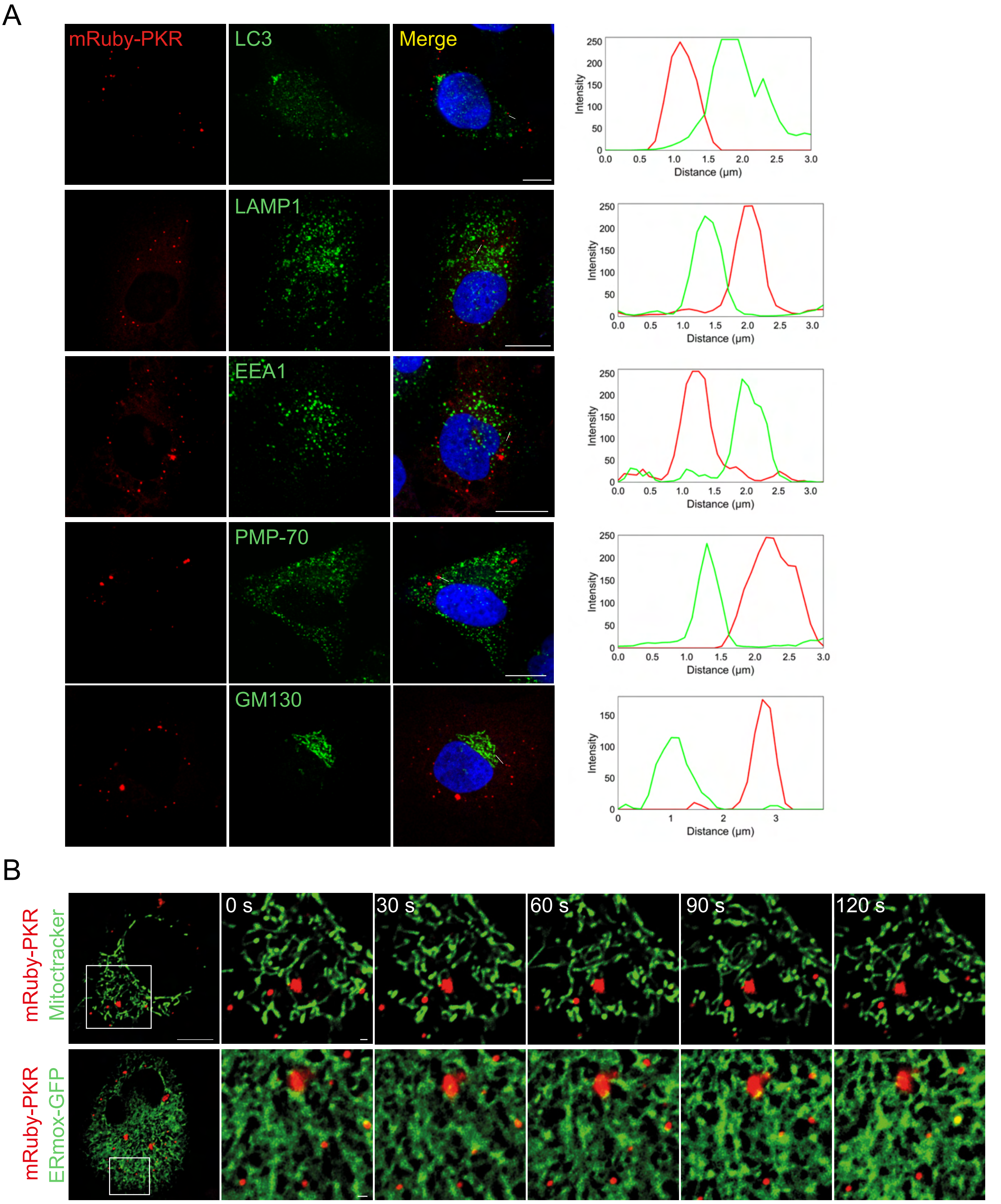

### Supplemental Figure 3

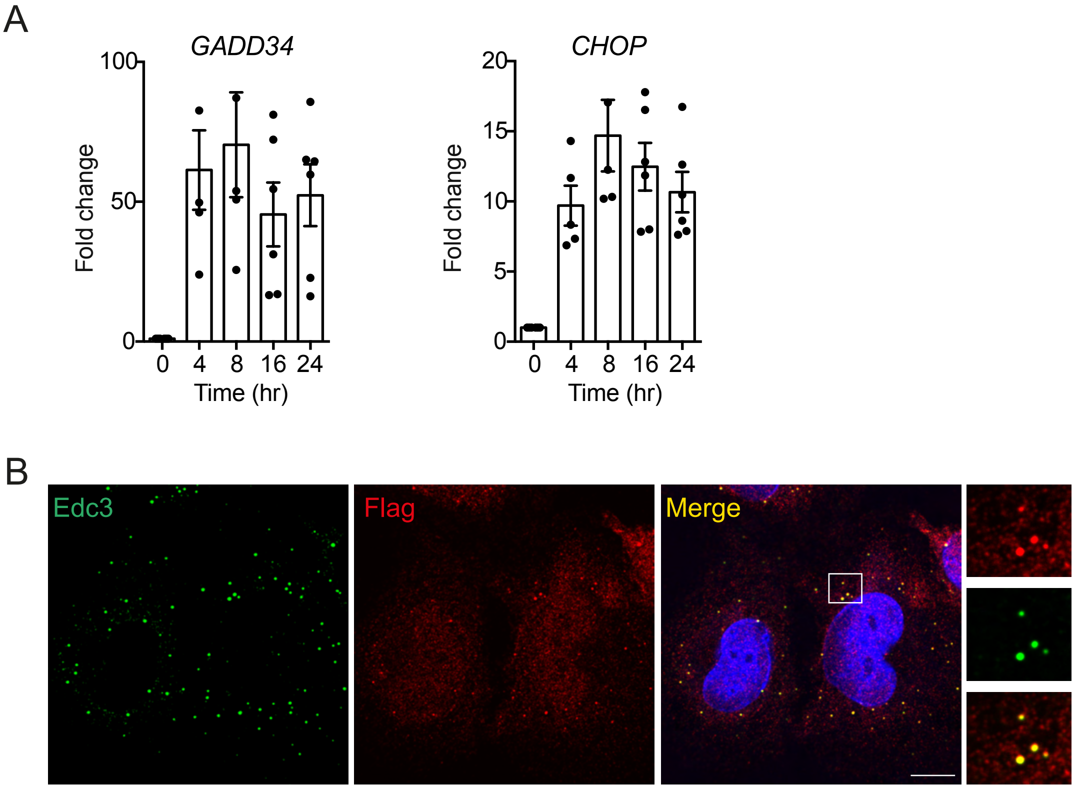

### Supplemental Figure 5

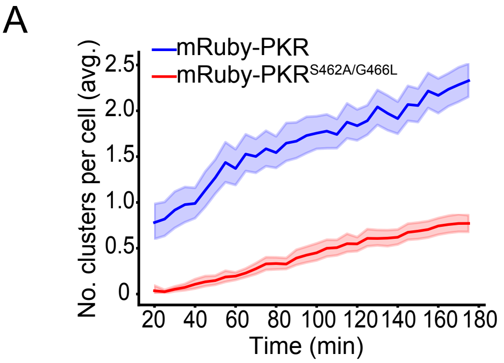

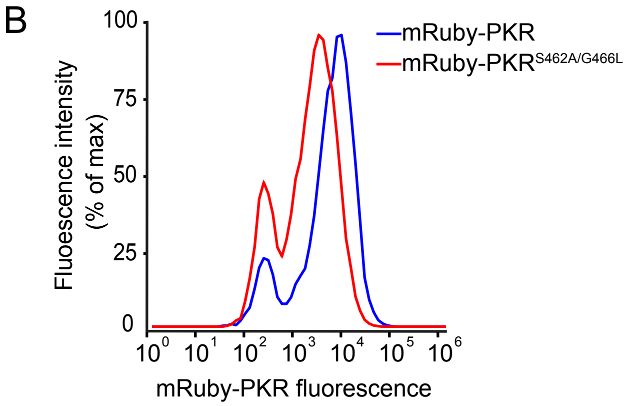

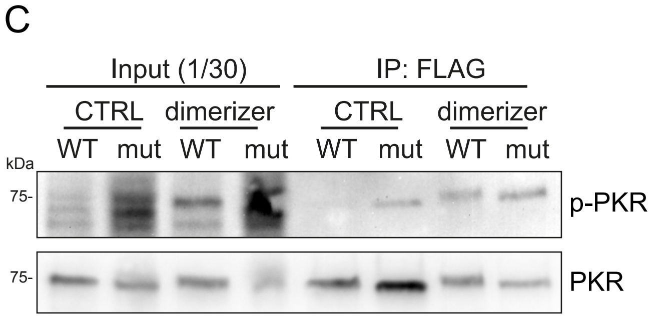
